# Patient-Derived Three-Dimensional Cortical Neurospheres to Model Parkinson’s Disease

**DOI:** 10.1101/2021.08.21.457201

**Authors:** Waseem K. Raja, Esther Neves, Christopher Burke, Xin Jiang, Ping Xu, Kenneth J Rhodes, Vikram Khurana, Robert H Scannevin, Chee Yeun Chung

## Abstract

There are currently no preventive or disease-modifying therapies for Parkinson’s Disease (PD). Failures in clinical trials necessitate a re-evaluation of existing pre-clinical models in order to adopt systems that better recapitulate underlying disease mechanisms and better predict clinical outcomes. In recent years, models utilizing patient-derived induced pluripotent stem cells (iPSCs) have emerged as attractive models to recapitulate disease-relevant neuropathology *in vitro* without exogenous overexpression of disease-related pathologic proteins. Here, we utilized iPSCs derived from patients with early-onset PD and dementia phenotypes that harbored either a point mutation (A53T) or multiplication at the α-synuclein/*SNCA* gene locus. We generated a three-dimensional (3D) cortical neurosphere culture model to better mimic the tissue microenvironment of the brain. We extensively characterized the differentiation process using quantitative PCR, Western immunoblotting and immunofluorescence staining. Differentiation and aging of the neurospheres revealed alterations in fatty acid profiles and elevated total and pathogenic phospho-α-synuclein levels in both A53T and the triplication lines compared to their isogenic control lines. Furthermore, treatment of the neurospheres with a small molecule inhibitor of stearoyl CoA desaturase (SCD) attenuated the protein accumulation and aberrant fatty acid profile phenotypes. Our findings suggest that the 3D cortical neurosphere model is a useful tool to interrogate targets for PD and amenable to test small molecule therapeutics.

## Introduction

More than 7 million Americans are suffering from neurodegenerative diseases, including Alzheimer’s Disease, Parkinson’s disease (PD), and Amyotrophic Lateral Sclerosis (ALS) (Ray Dorsey et al., 2018). There are no disease modifying therapies available for these disorders that can halt, slow or reverse disease progression. Predictive cellular models for human disease pathologies are critical for the development of effective therapeutics that may lead to successful clinical outcomes. Gene overexpression or knockdown models of neurological disease pathways in transformed or immortalized human cell lines have inherent limitations and often lead to unsuccessful outcomes in preclinical and clinical development (Horvath et al., 2016). In contrast, transgenic animal models of neurodegenerative disease can provide physiological and behavioral outputs, yet they are expensive, low-throughput, labor-intensive, and time-consuming. Most importantly, these models offer only limited insight into human disease mechanisms and potential therapeutic approaches (Dawson et al., 2018).

Induced pluripotent stem cells (iPSC) derived neuronal models provide a cellular model in which to study disease-relevant pathologies in patient neuronal cells. Integration of iPSC technology with genome editing to generate an isogenic control makes this approach unique and attractive, as disease-relevant mutations can be studied at endogenous expression levels in neuronal cells (Mungenast et al., 2019; Takahashi et al., 2007). In addition, iPSCs have unlimited renewal capabilities and can be differentiated into multiple neuronal-lineages, thereby providing a robust source of human neural cultures. Such a patient-derived cellular model may provide personalized drug-screening platforms that can be used to develop phenotypic assays and validate novel therapeutic targets in specific patients and test their efficacy (Zeltner and Studer, 2015). Using patient derived cells as a starting point also could allow for an *in vitro* personalized medicine approach, where the therapeutic effects of drugs can be tested at the individual patient cell level to identify and enrich for responders to particular treatments.

Combining iPSC technologies with three-dimensional (3D) cultures provides a multicellular spatial architecture to better approximate the native environment of the brain. There are two basic approaches to creating 3D culture systems either using a tissue engineering approach in which the neural cells are cultured in appropriate biocompatible 3D scaffolds such as Matrigel, or to culture aggregated stem cells as a suspension culture and differentiate them as organoids or spheroids, also known as neurospheres (Hopkins et al., 2015; Raja et al., 2016). Neurospheres can be generated from pre-differentiated neuronal stem cells (NSC) to reduce the duration of neuron maturation. The use of NSCs as a starting point provides an advantage as the cells are pre-defined to neuronal fate and it is possible to generate a consistent culture with lower variability in cellular composition.

The accumulation of pathogenic forms of α-synuclein (α-syn) is one of the primary pathological hallmarks of PD and the phosphorylation of serine 129 of α-syn (pS129 α- Syn) has been linked to PD and is enriched in Lewy bodies (LBs) (Oueslati, 2016). A recent study suggests that LBs are crowded with not just aggregated proteins but with lipid membranes including vesicular structures and dysmorphic organelles (Shahmoradian et al., 2019). The link between PD and the dysregulation of membrane lipids is growing stronger (Fanning et al., 2020). Studies have shown that dysregulation of monounsaturated fatty acids (MUFAs) can contribute to α-syn accumulation, while limiting MUFAs appears beneficial to α-syn-related pathology (Dettmer et al., 2015; Imberdis et al., 2019). Stearoyl CoA Desaturase (SCD) introduces a double bond into 16- and 18-carbon fatty acyl-CoA molecules (C16:0, palmityl CoA and C18:0, stearoyl CoA) to produce MUFAs (C16:1n7, palmitoleic acid and C18:1n9, oleic acid) that are incorporated into diverse lipid species, such as phospholipids, triacylglycerides, or cholesterol esters (Paton and Ntambi, 2009). Recent work from our group as well as others strongly suggests that targeting this pathway through SCD inhibition may prove to be therapeutic in alleviating several PD-related phenotypes seen in model systems (Fanning et al., 2018; Nuber et al., 2021; Vincent et al., 2018).

Here, we described the development of cortical neurosphere model from PD patients-derived iPSCs either carrying the A53T mutation in the gene encoding α-syn, *SNCA* (α- syn A53T) or triplication of the *SNCA* locus (S3) paired with corresponding mutation-corrected isogenic control. Compared to their isogenic controls, neurospheres from the PD patients showed dysregulated fatty acid profiles and accumulated total and phosphorylated α-syn levels. These disease-relevant phenotypes were reversed by a stearoyl-CoA desaturase inhibitor, CAY10566.

## Results

### Generation and Characterization of neural stem cells (NSC)

The iPSCs were carefully maintained by manually removing spontaneously differentiated cells. The A53T iPSC line and an isogenic control line (Corr) in which the A53T mutation was corrected by CRISPR-Cas9 genome engineering, were differentiated into neural stem cells (NSCs). After differentiation and purification (explained in the Materials and Methods section), initial quality control measures were implemented on both lines to assess the quality and purity of NSC. The NSC were banked and thawed for future experiments to generate two-dimensional or three-dimensional cultures. **Figure 1A** shows the entire process flow of the patient-derived *in vitro* model system utilized here. Phase contrast images of NSC showed similar morphology in both cell lines. Immunofluorescence microscopy also revealed similar staining patterns and levels of cell markers in both lines. Purified NSCs were positive for neural stem cell markers such as PAX6 and Nestin (**Figure 1B**). Additionally, the neural crest cell markers HNK1 and SOX10 were undetectable at this stage (**Figure 1C)**. Quantitative RNA analysis confirmed the immunostaining results; there were elevated levels of forebrain progenitor markers PAX6 in all NSC lines along with neural progenitor marker NESTIN and undetectable levels of neural crest cell marker Sox2. We also detected the presence MAP2 mRNA due to spontaneous differentiation of NSC into neurons (**Figure 1D)**.

**Figure 1.**
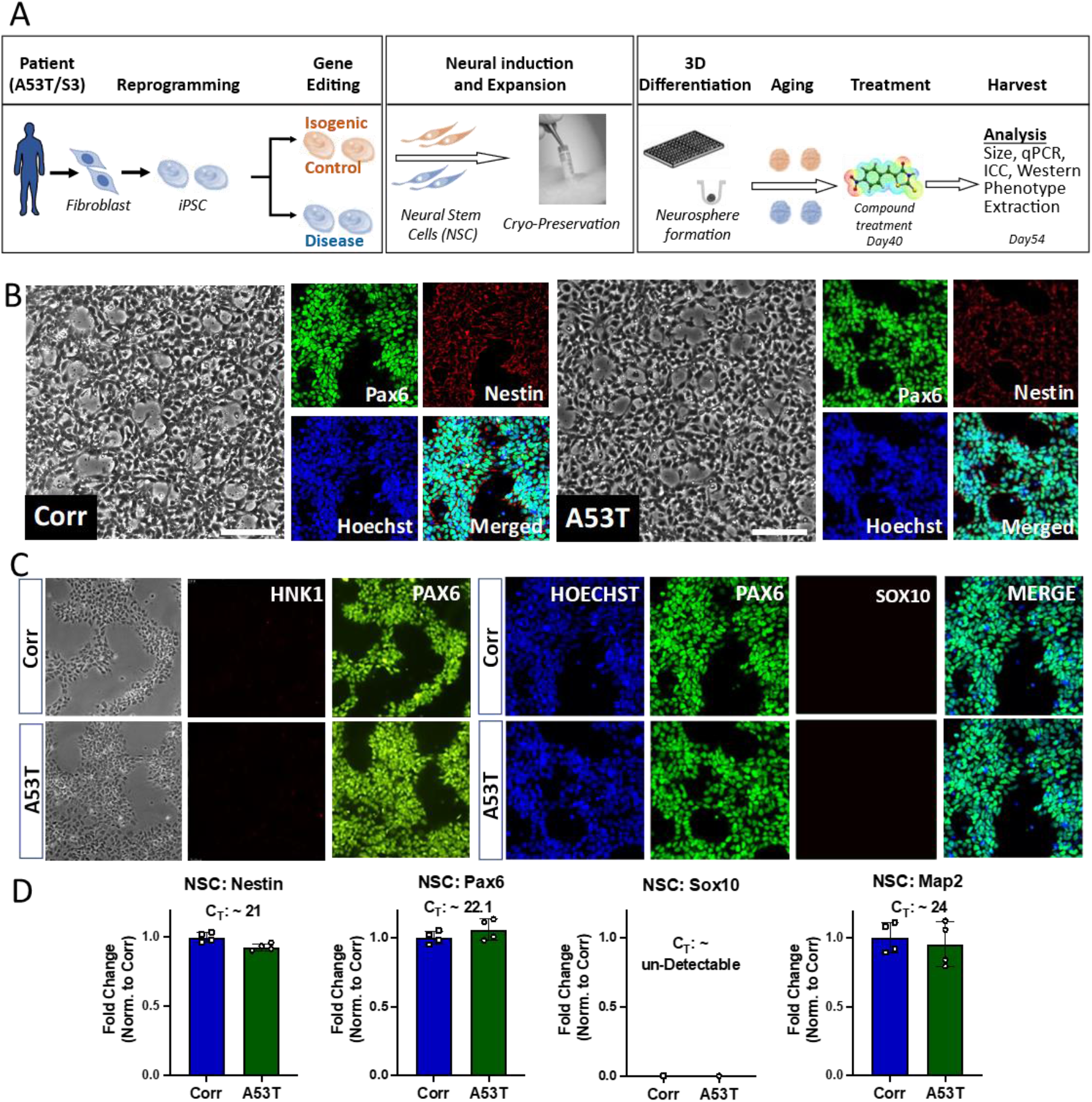
Process flow and Quality control of Neural Stem Cells (NSC). **A)** Process flow of patient-derived neurosphere generation. **B)** Representative phase contrast images and immunocytochemistry (ICC) for Hoechst and NSC markers (Nestin and PAX6). Left Panel: isogenic control line (Corr). Right panel: A53T line. **C)** ICC for Hoechst, PAX6 and neural crest cell markers (HNK1 and Sox10). Top panel: isogenic control line (Corr). Bottom panel: A53T line. **D)** Gene expression mRNA levels of PAX6, Nestin, SOX10 and MAP2 quantified using qPCR, with fold changes measured relative to the isogenic control. Scale bar: 500 µm.

The ability of neural stem cells to differentiate into neurons was then assessed as one of the quality controls measures. NSCs (A53T and Corr) were plated on a PDL/Laminin coated surface and differentiated in neural differentiation media (**Supplementary Table 1**) in a 2D format. After differentiation for 14 days, a clear neuronal morphology was observed with positive staining for the neuronal marker Tau as well as dendritic marker MAP2, in both lines (**Supplementary Figure 1**). These results together established that both the A53T and the isogenic control line (Corr) were capable of generating neural stem cells and neurons.

We also generated NSCs from patient-derived iPSCs harboring a triplication of the *SNCA* locus (S3) using the same protocol, along with the CRISPR-Cas9 generated isogenic control α-syn knockdown line (KD), where two copies of the *SNCA* genes were deleted to reduce the α-syn levels by ∼50%. The NSCs created from the S3 and KD iPSCs were evaluated using the same quality control measures as described above. Phase contrast imaging showed similar morphology between the S3 and KD NSCs (**Supplementary Figure 2A)**. Quantitative RNA analysis also confirmed positive expression of PAX6 and Nestin, and no expression of SOX10 was observed (**Supplementary Figure 2B)**. Additionally, NSCs derived from the S3 and KD lines successfully generated neurons in a monolayer differentiation culture (**Supplementary Figure 2C)**.

### Generation and Characterization of neurospheres

Three-dimensional (3D) neurosphere cultures may provide a more relevant and complex environment to better mimic brain tissue. A protocol for generating neurospheres was therefore optimized for the A53T/Corr and the S3/KD neural stem cells. The neurospheres were formed in an ultra-low adhesion 384 well format. It was noted during the differentiation and maturation process that sphere size gradually increased based on quantitation of phase contrast images (**Figure 2A, Supplementary Figure 3A**). RNA levels of relevant marker genes were then further investigated, and no significant difference was observed in levels of MAP2, Synapsin-I, S100B (astrocyte marker) and Pax6 in patient versus isogenic control neurospheres, although the Pax6 level was drastically reduced compared to NSCs (**Figure 2D, Supplementary Figure 3D)**. Immunofluorescence microscopy for MAP2 and S100B corroborated the RNA analyses for the A53T/Corr and S3/KD neurospheres (**Figure 2B, Supplementary Figure 3B**). The glutamatergic neuron marker VGLUT2 was abundantly expressed in these cultures whereas GABAergic neuron marker GAD1/67 expression was largely absent (**Figure 2C, Supplementary Figure 3C)**. The GAD1/67 antibody was validated using iCell GABA neurons (Fujifilm) as a positive control and iCell Gluta neurons (Fujifilm) as a negative control (**Supplementary Figure 3E)**. These results suggest that glutamatergic neurons are the major neuronal population in the cortical neuropheres.

**Figure 2.**
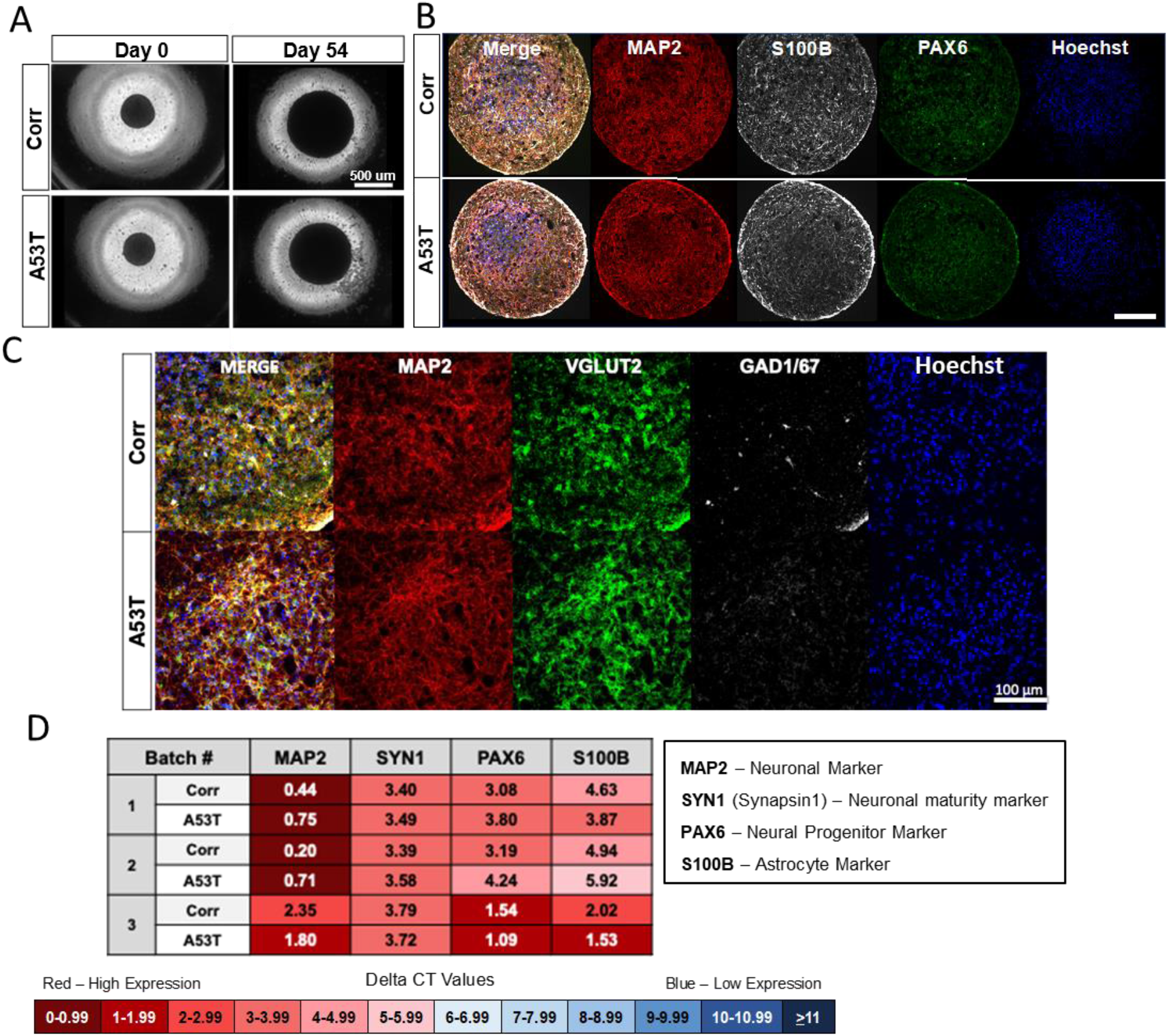
Quality control for A53T/Corr neurosphere differentiations shows cellular subtype composition of cultures. **A)** Representative phase contrast images of neurospheres at day 0 and day 54. Top Panel: Isogenic control line (Corr). Bottom panel: A53T line. **B)** Immunocytochemistry (ICC) of 54-day-neurospheres for neuronal marker MAP2, astrocyte marker S100B, NSC marker PAX6, and Hoechst. Top panel: Isogenic control line (Corr). Bottom panel: A53T line. Scale bar: 200 µm. **C)** ICC of 54-day-neurospheres for MAP2, glutamatergic neuron marker VGLUT2, GABAergic neuron marker GAD1/67 and Hoechst. Top panel: Isogenic control line (Corr). Bottom panel: A53T line. Scale bar: 100 µm. **D)** Gene expression of MAP2, PAX6, S100B, and Synapsin-I in 54-day-neurospheres for three representative neurosphere differentiations. Delta CT values, measured by qPCR, are the difference between the CT values of the housekeeping gene (GAPDH) and the gene of interest. A lower CT value (in red) indicates high gene expression, while a high CT value (in blue) indicates low gene expression.

### Phenotypic assessment in NSC and neurospheres

#### Neurospheres from patient-derived iPSC display an aberrant fatty acid profile

Recent work from our group and others identified SCD as potential targets to ameliorate α-syn toxicity (Fanning et al., 2018; Nuber et al., 2021; Vincent et al., 2018). These studies implicate fatty acid biology in mediating aspects of α-syn toxicity, which prompted the evaluation of fatty acid profiles in neural stem cells and mature neurospheres derived from A53T/ Corr and S3/KD lines.

26 saturated and unsaturated fatty acids ranging from 14 to 22 carbon chain lengths were profiled in NSCs from all lines, and little significant differences were observed between the disease lines (A53T and S3) and corresponding isogenic control lines (Corr and KD), with the exception of an elevated C16 desaturation index in the A53T NSCs (**Figure 3A and 3B**). The C16 desaturation index was calculated as the ratio of palmitoleic acid (C16:1n7) to palmitic acid (C16:0). In neurospheres cultured for 54 days, there were several changes consistent between the disease (A53T and S3) lines and isogenic control lines. These included significant increases in both the C16 and C18 (C18:1n9/C18:0) desaturation indices in the A53T and S3 lines compared to the respective isogenic control lines **(Figure 3C and 3D)**. The level of an elongation product of C16:1n7, C18:1n7 was also significantly elevated in the disease lines compared to the control lines (**Supplementary Figure 4 and 5**). In contrast, an essential fatty acid such as linoleic acid (C18:2n6) and polyunsaturated fatty acids (PUFA), such as γ-linolenic acid (C18:3n6) and docosapentaenoic acid (C22:5n3), were significantly reduced in the disease (A53T and S3) neurospheres as compared to the respective isogenic control (Corr and KD) neurospheres (**Figure 3C and 3D, Supplementary Figure 4 and 5**). These data suggest that the PD mutations including A53T α-synuclein mutation or elevated levels of wild type α-synuclein in the SNCA triplication influences fatty acid homeostasis in patient-derived neurons.

**Figure 3.**
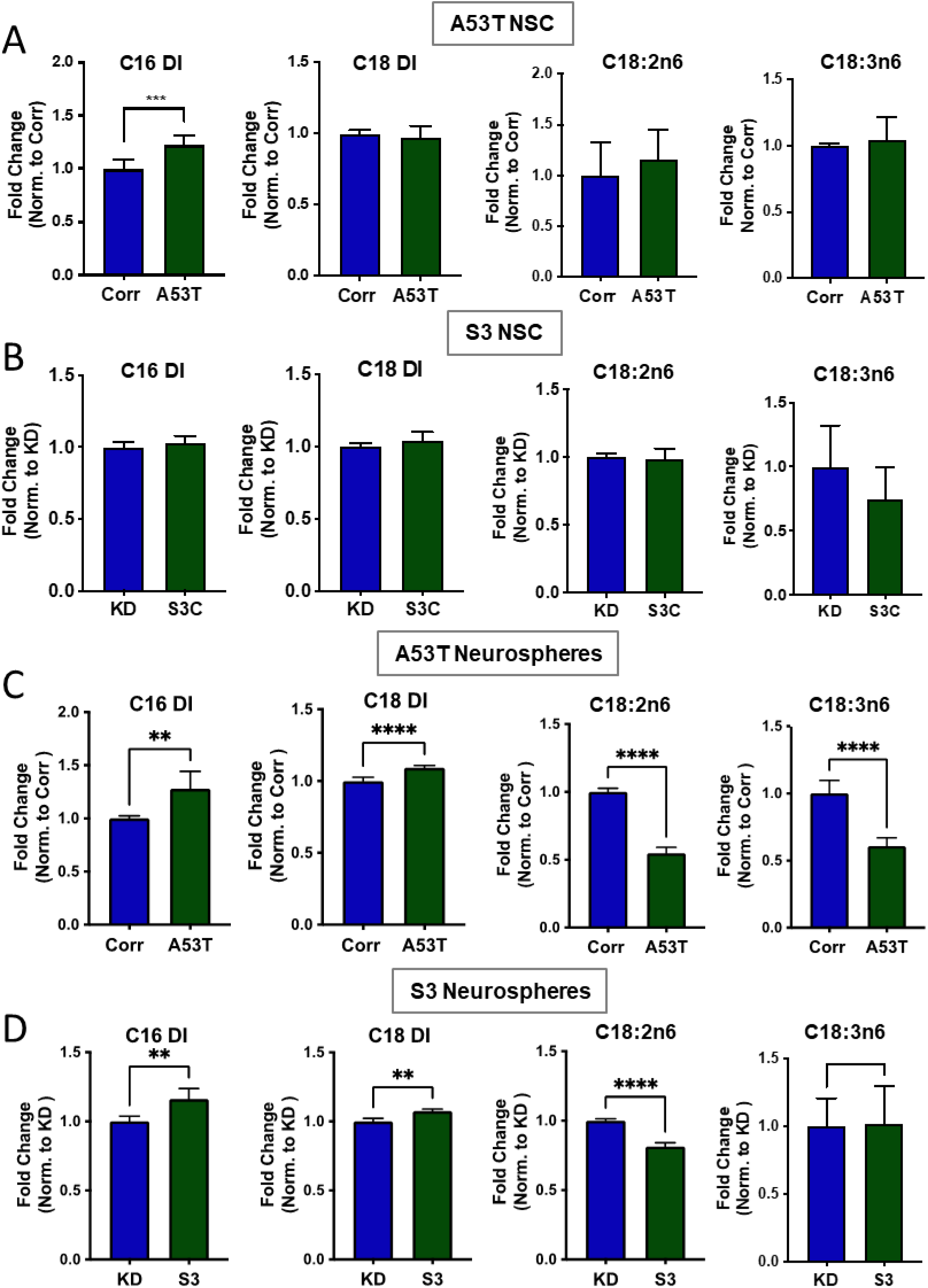
Fatty acid profile of NSC and neurospheres. Desaturation index (DI) for C16 (C16:1n7/C16:0) and C18 (C18:1n9/C18:0) along with the relative levels of the essential fatty acids, linoleic (C18:2n6) and γ-linolenic acid (C18:3n6) for the A53T/Corr **(A**) and S3/KD **(B)** NSC and the A53T/Corr **(C**) and S3/KD **(D)** neurospheres at day 54. The analysis was conducted using an unpaired two-tail t-test, with n=6 for **(A)** and n=5 for **(B)**. The analysis for neurospheres was conducted using an unpaired t-test, with data from three different neurosphere differentiations. Each differentiation has two biological replicates. Data are mean ± SD and analyzed by two-tail t-test. *P<0.05; **P≤0.01; ***P≤0.005; ****P≤0.0001

#### Patient-derived neurospheres display an age-dependent increase in total and phospho-serine129 α-synuclein

Increased levels of α-syn are linked to sporadic and familial PD as well as Lewy body dementia and multiple system atrophy (Mollenhauer et al., 2019). In addition, phosphorylation of serine residue 129 (pS129) in α-syn has been linked to PD and is enriched in Lewy bodies (Fujiwara et al., 2002), causing it to gain traction as biomarker for the disease. Both α-syn and pS129 α-syn levels were measured in the NSC and the cortical neurospheres from these two pairs of PD patient lines using Western blot. At the NSC stage, pS129 α-syn was not detected, while a modest increase in total α-syn levels was observed in the A53T line compared to the Corr line, however the S3 line showed significantly elevated levels of total α-syn compared to the KD line **(Figure 4A and 4B)**. After undergoing cortical neuron differentiation and 54-days of maturation, the neurosphere cultures displayed increased levels of total α-syn protein and a more dramatic increase in pS129 α-syn in the A53T neurospheres compared to the Corr control (**Figure 4C and 4E**). This phenotype was consistent throughout multiple rounds of differentiation. Additionally, we observed a similar phenotype in the S3 triplication line where the S3 neurospheres displayed increase in both total α-syn and pS129 α-syn compared to the KD control neurospheres (**Figure 4D and 4F**).

**Figure 4.**
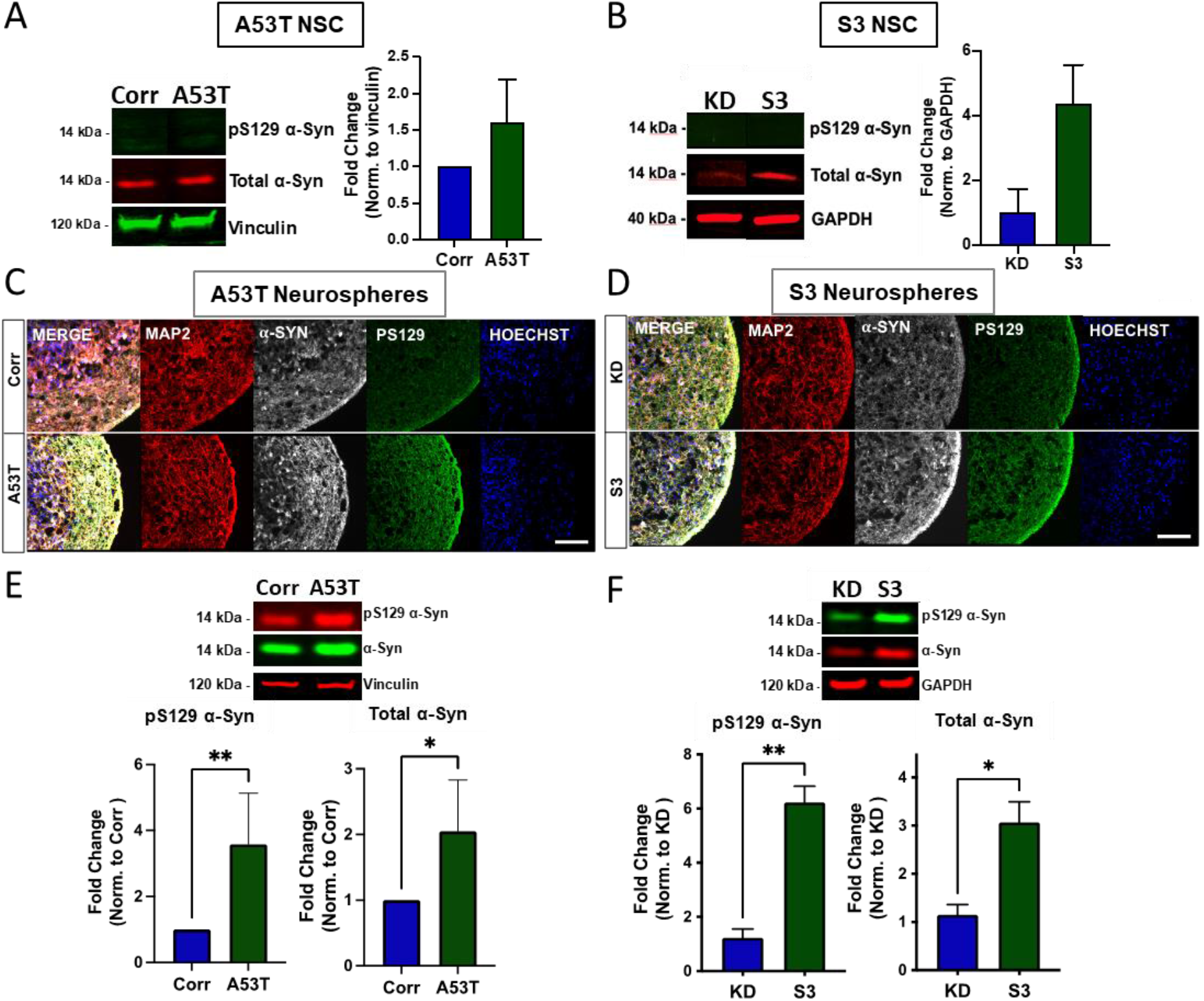
Total and pS129 α-synuclein in NSC and neurospheres. **A and B)** Representative Western blot for pS129 and total α-synuclein and quantification for total α-synuclein in the A53T/Corr (**A**) and S3/KD NSC (**B**). **C and D)** Immunostaining of A53T/Corr (**C**) and S3/KD (**D**) neurospheres at day 54 for MAP2, α-synuclein, pS129, and Hoechst. Scale bar: 100 µm. **E and F)** Representative Western blot (top panel) and quantification (bottom panel) for pS129 and total α-synuclein for A53T/Corr (**E**) and S3/KD (**F**) neurospheres at Day 54. The quantification data is an aggregate of five separate differentiations for the A53T/Corr lines, and two differentiations for the S3/KD lines. Data are mean ± SD and analyzed by two-tail t-test. *P<0.05; **P≤0.01

### Effects of SCD inhibitor CAY10566 on A53T/Corr and S3/KD neurospheres

We next treated 40-day-old neurospheres with a potent SCD inhibitor, CAY10566 to see if SCD inhibition can correct the change in fatty acid profiles in the disease lines. Treatment of A53T and S3 neurospheres with 0.3 μM CAY10566 for 14 days reversed the elevation of C16 and C18 DI in both lines (**Figure 5A and B**). Interestingly, the reduced levels of linoleic (C18:2n6) and γ-linolenic acids (C18:3n6) in the A53T line and linoleic acid (C18:2n6) in the S3 line were also reversed by CAY10566 (**Figure 5A**). A similar effect of CAY10566 was also observed in the isogenic control lines, however the magnitude of change was not as robust as observed in the disease lines (**Figure 5A and B**). These results suggest that SCD inhibition alleviated the abnormal fatty acid profiles in PD patient-derived neurospheres.

**Figure 5.**
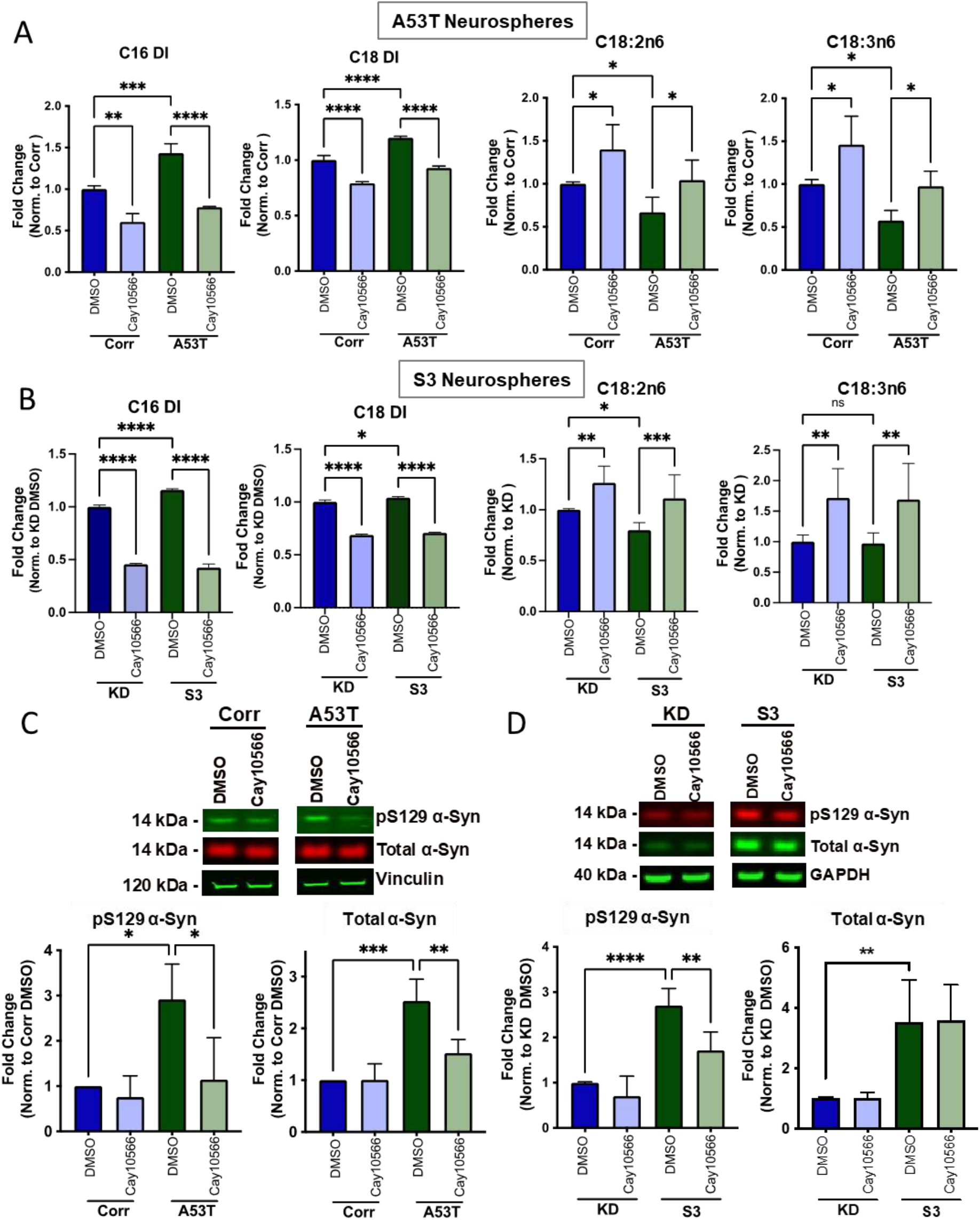
Treatment with the SCD inhibitor CAY10566 reversed the abnormal fatty acid profiles and pathogenic α-synuclein levels. **A** and **B**) C16 and C18 desaturation index (DI) and essential fatty acids Linoleic (C18:2n6) and γ-Linolenic acid (C18:3n6) from A53T/Corr (**A**) and S3/KD (**B**) neurospheres treated for 2 weeks with 0.3 μM CAY10566. Data are from three independent neurosphere differentiations. All values in each differentiation were normalized to the isogenic control treated with DMSO in each respective batch of differentiation. **C and D)** Representative Western blot (top panel) and quantification (bottom panel) for pS129 and total α-synuclein in 54 day-old cortical neurospheres from A53T/Corr (**C**) and S3/KD (**D**) treated for 2 weeks with 0.3μM CAY10566. Quantification includes data from three (A53T) and two (S3) distinct differentiations, with two biological replicates per group. Data are mean ± SD and analyzed by One-way ANOVA analysis with Tukey’s multiple comparison test. *P<0.05; **P≤0.01; ***P≤0.005; ****P≤0.0001

The same treatment paradigm with CAY10566 also reduced total and pS129 α-syn levels in A53T neurospheres (**Figure 5C**). CAY10566 also resulted in a similar reduction of pS129 α-syn in the S3 lines (**Figure 5D**), while total α-syn levels were not greatly impacted in the S3/KD line. These results suggest that SCD inhibition attenuated the abnormal accumulation of α-syn in PD patient-derived neurospheres.

## Discussion

Parkinson’s disease (PD) is the second most common irreversible neurodegenerative disease today. Though the disease was first described over 200 years ago, there is no effective therapy to stop, slow, or reverse the progression of PD. The first step to tackle this problem is to develop predictive models that help us to recapitulate and understand the pathophysiology of the disease. This recapitulation of the PD pathophysiology can then be harnessed to test the effects of therapeutic compound treatment.

Post-mortem human brain tissue from PD patients is an invaluable resource that helped us to identify key mutations in various genes including α-synuclein that lead to PD (Klein and Westenberger, 2012). However, it only offers the analysis at the end stage of the disease and is not amenable to drug manipulation or screening therapeutic agents. Engineered rodent models based on disease-linked mutations have helped understand the progression of pathology in animals, but the intrinsic differences between animals and humans may contribute to the lack of translation in efficacy from animals to the clinic (Dawson et al., 2018). Considering these limitations, using patient-derived cellular models generated by induced pluripotent stem cells (iPSC) may provide an attractive alternatives approach (Mungenast et al., 2019; Zeltner and Studer, 2015).

Different groups have explored the implications of various α-synuclein gene mutations in iPSC derived neural cells, such as the A53T point mutation and SNCA copy number variants triplication mutation. In 2013, Chung and colleagues utilized the A53T patient line to show the phenotypic differences between the A53T and isogenic control lines (Chung et al., 2013). In 2014, Flierl and colleagues investigated the vulnerabilities of NSC carrying an SNCA triplication mutation (Flierl et al., 2014). In our own exploration of the α-syn A53T and triplication mutations (S3), we utilized a non-integrated mRNA-based reprogramming strategy to generate our iPSC from the PD patients followed by CRISPR-Cas9 gene editing technique to create isogenic control lines (Corr and KD).

Patient-derived iPSC can be instructed towards an ectoderm lineage and differentiated into neural stem cells. Dual-SMAD inhibition (BMP and TGF-β signaling inhibition) is the most common technique used to generate NSC from iPSC, and number of published literatures has shown the utility and efficiency of this technique both 2D (monolayer) and 3D (Embryoid body) differentiations. These discoveries led to the development of commercial kits to generate NSC from iPSC and we utilize one such commercial kit (Scientific) in our study to differentiate iPSC into NSC in a monolayer format. We performed extensive characterization of our NSC, and we have not seen any differences (at the mRNA and protein levels) between the disease and isogenic control lines.

We further differentiated NSC into neurons and astrocytes in a 3D suspension culture format without the use of any supporting biomaterial. One of the biggest advantages of the 3D neurospheres generated from these NSC is its reproducibility by starting from well characterized NSC and its low maintenance. Additionally, in the 3D format, the ease of compound treatment and scalability of the cultures allows more cellular material to undergo high throughput screening. In addition, 3D cultures can be aged more easily than 2D cultures, which tend to lift off from the coated cell culture surface during aging. This is important when aging is an important factor in the emergence of disease relevant phenotypes. Our results demonstrated using the accumulation of α-syn and the disruption of fatty acid profiles in the aged neurospheres but not in NSC.

Although neurospheres provide a more accessible platform to age our neurons, they need careful monitoring and considerable quality control measures to determine the differentiation quality. Since the starting cells for the differentiation are NSC, the resulting cells are limited to a neural or astrocytic lineage. A typical differentiation yields in mostly glutamatergic neurons that are VGLUT2- and MAP2-positive, with some GABAergic neurons that are GABA-positive with the presence of S100β-positive astrocytes. If the disease and control lines exhibit substantially different cellular subtype compositions, it can affect the downstream analyses and compound treatments. In that aspect, it is crucial to monitor the composition of cellular subtypes to ascertain that any phenotypes observed are due to genotypic differences, not differentiation differences between the lines. Overall, the neurosphere differentiation method yields cell cultures with consistent quality and robust disease-relevant phenotypes.

While still an emerging field, the link between Parkinson’s Disease and disrupted lipid biology is becoming stronger (Fanning et al., 2020). Our results add to this growing evidence by identifying elevation of C16 and C18 DI along with other altered fatty acids in the A53T and S3 patient-derived neurospheres. This phenotype appears to be age-dependent, as no significant changes are observed at the NSC stage. Moreover, this aberrant lipid desaturation phenotype appears in both our A53T mutant and SNCA triplication lines, suggesting that dysregulation of lipids and altered fatty acid saturation may be a key pathological event occurring in synucleinopathies and common across different genetic forms of the disease.

The accumulation of pathogenic forms of α-syn is one of the primary pathological hallmarks of PD and the phosphorylation of serine 129 of α-syn has been linked to PD and is enriched in LBs (Oueslati, 2016). These findings have been recapitulated in several models of PD, ranging from human α-syn overexpression models to iPSC neurons derived from PD patients harboring mutations in α-syn or the PD risk factor (Ludtmann et al., 2018; Mazzulli et al., 2016; Prots et al., 2018; Ryan et al., 2013). Here, we have shown the age-dependent accumulation of total and pS129 α-syn in PD patient-derived cortical neurospheres, consistent with previous studies across PD models. Interestingly, we observe that two-week treatment of SCD inhibition with CAY10566 not only reversed the aberrant fatty acid profiles but also reduces the accumulation of pS129 α-syn in aged neurospheres from both the A53T and the S3 patients. The consistent findings in two patient lines raise an intriguing possibility that SCD inhibition may prove to be therapeutic in alleviating several PD-related phenotypes as seen in model systems (Fanning et al., 2018; Nuber et al., 2021; Vincent et al., 2018).

Taken together, our cortical neurosphere protocol provides a useful and robust tool to develop therapeutics against devastating neurodegenerative diseases by enabling identification of age-dependent disease-relevant phenotypes and testing therapeutic agents in patient-derived cells.

## Materials and Methods

### iPSC generation, Editing and Maintenance

All cells were maintained in the incubator at 37°C with 5% CO_2_. The A53T patient-derived iPSCs were generated from fibroblasts obtained from a patient skin biopsy at Boston University. The fibroblast to iPSC reprogramming was performed at the Harvard Stem Cell Institute iPS Core using a non-integrative (mRNA) strategy. The α-Syn triplication patient cell line (S3) was purchased from the Coriell Institute for Medical Research. The iPSCs were expanded and maintained in mTeSR1 medium from Stem Cell Technologies (see iPSC Media in Supplementary Table 1).

### Generation of Neural Stem Cells (NSC)

The iPSC were differentiated into Neural Stem Cells using a monolayer method. Briefly, the iPSC were plated and kept in NSC induction media (see **Supplementary Table 1**) for seven days. On day seven, the cells were re-plated in NSC induction media, and switched to NSC expansion media (NEM, see **Supplementary Table 1**) the next day. Neural crest cells were removed by Accutase treatment and passaging (see supplementary data for detail). After achieving a pure population of NSC, the NSC were expanded and frozen in Synth-a-freeze Cryopreservation Medium (**Figure 1A**).

### Differentiation of NSC into neurons in 2D

NSC were maintained and expanded in NEM. For 2D neuron differentiation, the NSC were plated in NEM. The next day, neuron differentiation media was added to the NSC (the same as neurosphere differentiation media in **Supplementary Table 1**). The cells were kept in this media for 10 to 12 days. On day 10-12, the neurons were terminally plated in the desired plate format. The neurons were aged for 2 to 4 weeks in neuron maturation media (the same as neurosphere maturation media in **Supplementary Table 1**). At the end of the experiment, the neurons were washed once with DPBS and fixed in 4% paraformaldehyde for immunocytochemistry.

### Generation of neurospheres

The NSC were passaged and plated as a single cell suspension in 384 well ultra-low adhesion spheroid plates at a density of 20,000 cells per well. After loading cells, the spheroids plates were centrifuged at 300g for 5 minutes to aggregate the cells and to remove air bubbles. The next day, the NEM was replaced with the neurosphere Differentiation Media (see **Supplementary Table 1**). The differentiation media was changed every three days. After ten days of differentiation, the neurosphere Differentiation Media was replaced by the neurosphere Maturation Media (see **Supplementary Table 1**). The media was changed bi-weekly for the next three weeks, and then once a week after day 30. At fixed harvesting times, the neurospheres were washed with DPBS and stored in -80°C (for RNA, Western Blot, or FADI analysis) or fixed in 4% paraformaldehyde for cryosectioning (**Figure 1A**).

### Compound treatment

40 day-old neurospheres from both patient-derived and isogenic control lines were treated with 0.3 uM CAY10566 along with DMSO as a vehicle. The final concentration of DMSO in the media was 0.03%. The spheres were treated twice a week and harvested after two weeks of treatment. The spheres were harvested and pooled from multiple wells of the 384 well plates for different assays.

### RT-qPCR

NSC, and neurospheres were processed for RNA extraction using the RNeasy Plus Mini Kit (from Qiagen). The extracted RNA was converted to cDNA with the qScript cDNA Supermix (from Quanta Biosciences) and the Mastercycler ep using RT-PCR. The cDNA was loaded in a 96 well or 384 well qPCR plate at a starting RNA concentration of 20 ng/ul for all the samples. Gene expression was then analyzed with qPCR using the Taqman Fast Advanced Master Mix (Thermofisher), Taqman Gene Expression Probes, and the Step One Plus Real-Time PCR System. The list of genes run for qPCR are in **Supplementary Table 4**.

### Immunocytochemistry

NSC and neurons were cultured in 96 well plates and fixed in 4% paraformaldehyde (PFA). The NSC were stained for different neuronal stem cell markers (PAX6 and Nestin), as well as the neural crest marker SOX10. Neurons were stained for different neuronal markers (MAP2, total Tau, VGLUT2 and GABA) and an astrocyte marker (S100B). The neurospheres were harvested as mentioned above and fixed in 4% PFA overnight. The neurospheres were cryosectioned (Histoserv Inc) and stained using a published protocol (Raja et al., 2016). The stained slides were imaged on a Nikon confocal microscope. The catalog number and sources of all the primary and secondary antibodies are listed in **Supplementary Table 2 and 3**.

### Western blotting

The harvested neurospheres stored at -80C were thawed and lysed according to a published protocol with modifications (ref) and protein concentrations were measured using the BCA method (Thermofisher). The lysates were run on 4-12% SDS PAGE gels (Invitrogen), and dry transferred to 0.22 uM PVDF membranes. The blots were stained with primary antibodies and Li-Cor secondary antibodies and were imaged on a Li-Cor Odyssey CLX. The catalog number and sources of all the primary and secondary antibodies are listed in **Supplementary Table 2 and 3**.

### Fatty Acid Desaturation Index (FADI) Analysis

The neurospheres were harvested and stored in 80% methanol at -80°C. The samples were shipped (on dry ice) to OmegaQuant LLC and processed for different fatty acids using gas chromatography (GC) with flame ionization detection. Total lipids were extracted from NSC and neurospheres and hydrolyzed, individual fatty acids were derivatized as methyl esters, and then assessed by gas chromatography with flame ionization detection. Fatty acid composition was expressed as a percent of total identified fatty acids. The abundance of different fatty acid was quantified using Microsoft Excel and plotted using GraphPad Prism software.

### Statistical Analysis

The results are presented as the mean ± standard deviation. All statistical analyses were performed using GraphPad Prism 7.0 software. The p values were calculated with a two-tailed Student’s t-test and ONE-way ANOVA (Tukey’s test *p < 0.05, **p < 0.01, ***p < 0.001). Results shown here are the representative data from greater than 5 experiments performed independently at different time points by multiple individuals.

