## Supplementary-Figures Materials and Methods for "Patient-Derived Three-Dimensional Cortical Neurospheres to Model Parkinson’s Disease"

### Title

### Authors and Affiliation

Waseem K. Raja[1*], Esther Neves[1], Christopher Burke[1], Xin Jiang[1], Ping Xu[1], Kenneth J Rhodes[1†], Vikram Khurana[2,3,4], Robert H Scannevin[1‡], Chee Yeun Chung[1*]

[1]Yumanity Therapeutics, Boston, MA, 02135, USA
[2]Ann Romney Center for Neurologic Disease, Department of Neurology, Brigham and Women's Hospital and Harvard Medical School, Boston, MA 02115, USA
[3]Harvard Stem Cell Institute, Cambridge, MA 02138, USA
[4]Broad Institute of MIT and Harvard, Cambridge, MA 02142, USA
[†]Current address: Wave Life Sciences, Cambridge MA, 02138, USA
[‡]Current address: Verge Genomics, South San Francisco, CA, 94080, USA
* Corresponding author

### Supplementary Figures

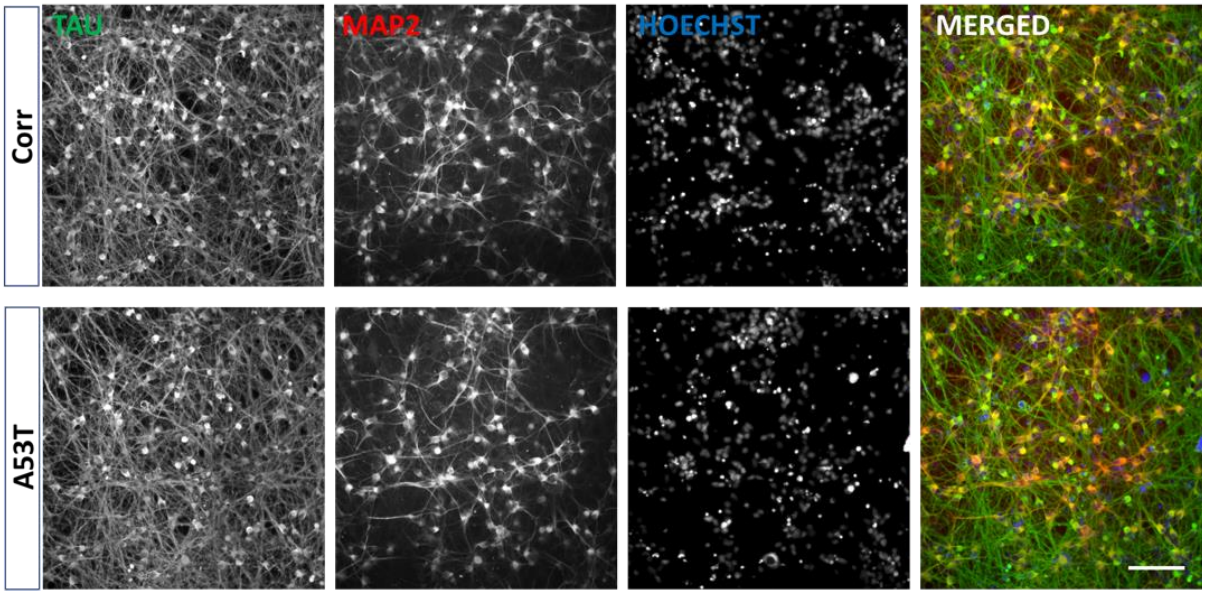


**Supplementary Figure 1. Representative Immunocytochemistry (ICC) stained images of neurons differentiated form disease and isogenic control.** The NSC were expanded, differentiated into neurons, terminally plated, and fixed after two weeks in culture. Top and bottom panel represent isogenic control and disease neuron, respectively. The neurons were stained with an anti-Tau antibody (green) for whole neuron staining, MAP2 (red) and Hoechst for nuclear stain (blue) Scale bar: 200 µm.

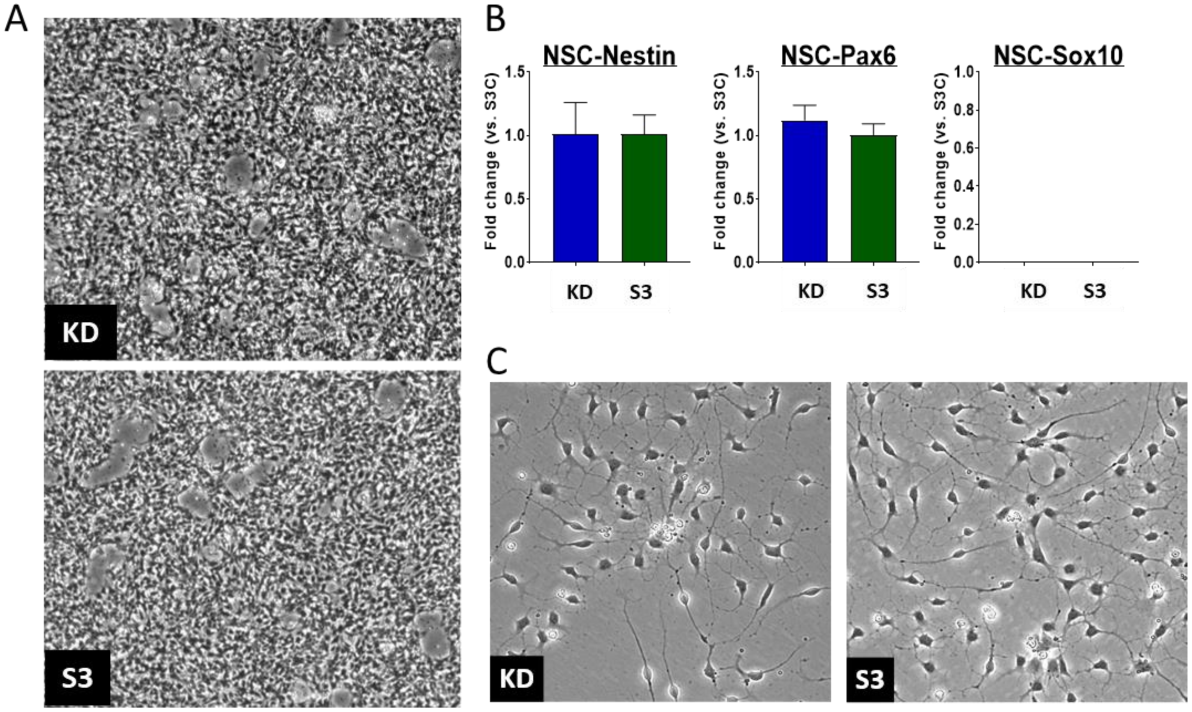


**Supplementary Figure 2**. **iPSC to Neural Stem Cell differentiation from S3 and KD lines. A)** Neural Stem cell phase contrast images from KD (top) and S3 (bottom). **B)** mRNA level was quantified using qPCR for S3 and KD lines and fold change were measured relative to S3 for NSC marker PAX6, Nestin along with Neural Crest Cell marker SOX10. **C)** Representative images of NSC differentiated into neuron in a monolayer culture from S3 and isogenic control lines. Scale bar: 500 µm.

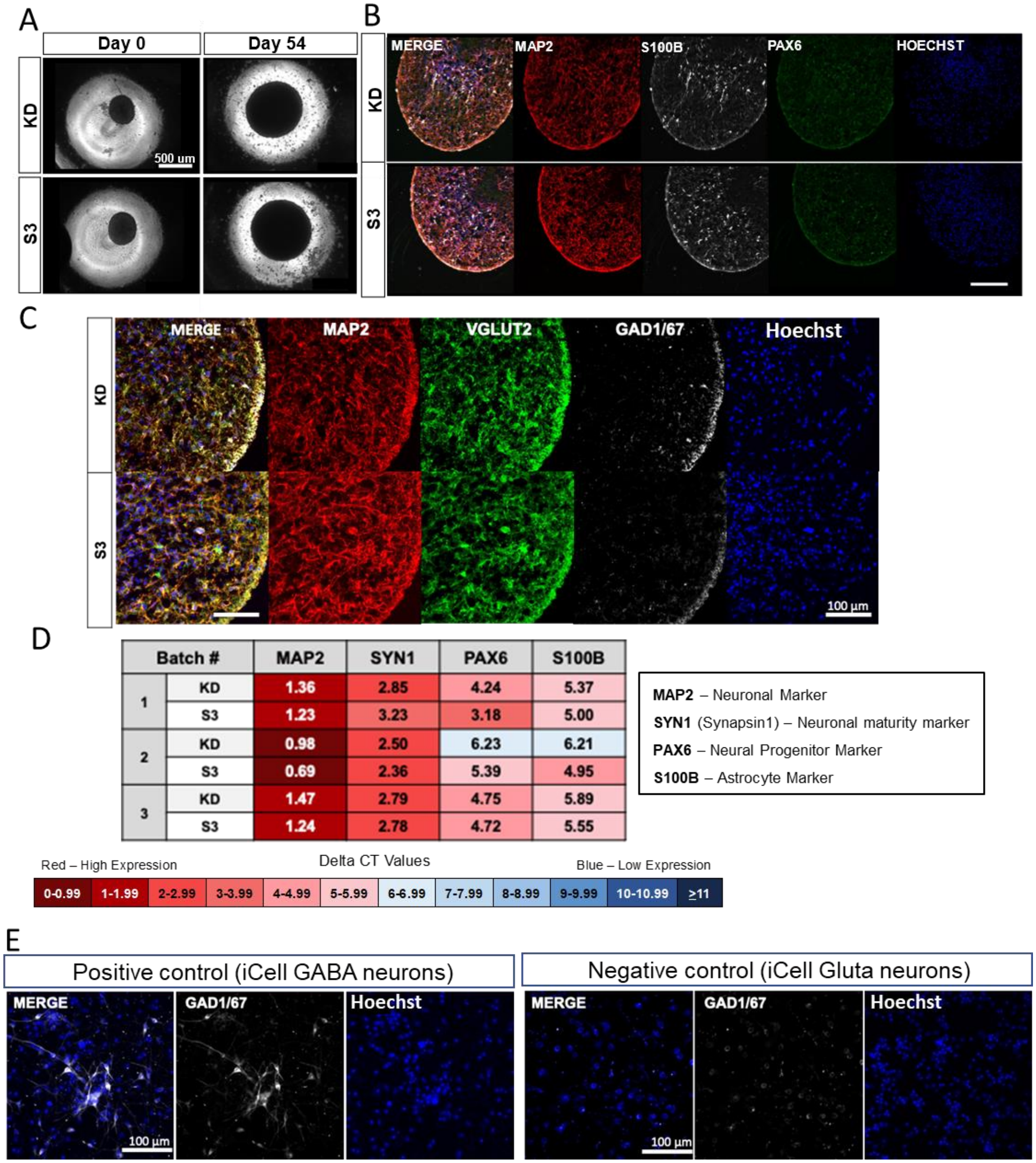


| Batch # | | MAP2 | SYN1 | PAX6 | S100B |
|---|---|---|---|---|---|
| 1 | KD | 1.36 | 2.85 | 4.24 | 5.37 |
| | S3 | 1.23 | 3.23 | 3.18 | 5.00 |
| 2 | KD | 0.98 | 2.50 | 6.23 | 6.21 |
| | S3 | 0.69 | 2.36 | 5.39 | 4.95 |
| 3 | KD | 1.47 | 2.79 | 4.75 | 5.89 |
| | S3 | 1.24 | 2.78 | 4.72 | 5.55 |

**Supplementary Figure 3 Quality control for S3/KD neurosphere differentiations shows cellular subtype composition of cultures. A)** Phase contrast images show that over the 54-day maturation period neurospheres grow from around 400 µm in diameter to 600 – 900 µm in diameter. **B)** Immunocytochemistry of cryosectioned neurospheres shows the expression of neuronal marker MAP2, astrocyte marker S100B, NSC marker PAX6, and nuclear stain Hoechst for S3 and KD neurospheres. **C)** Immunocytochemistry of cryosectioned neurospheres shows the expression of neuronal marker MAP2, glutamatergic neuron marker VGLUT2, GABAergic marker GAD1/67 and nuclear stain Hoechst in the S3 and KD neurospheres. **D)** The delta CT value for the gene of interest in S3/KD neurospheres. A lower CT value (in red) indicates high gene expression, while a high CT value (in blue) indicates low gene expression. Three representative batches of differentiation are shown for four genes of interest: MAP2 (Neuronal Marker), Synapsin-I (Neuronal Maturity Marker), PAX6 (NSC Marker) and S100B (Astrocyte Marker). **E)** The GAD1/67 antibody was validated using iCell GABA neurons (Fujifilm) as a positive control and iCell Gluta neurons (Fujifilm) as a negative control.

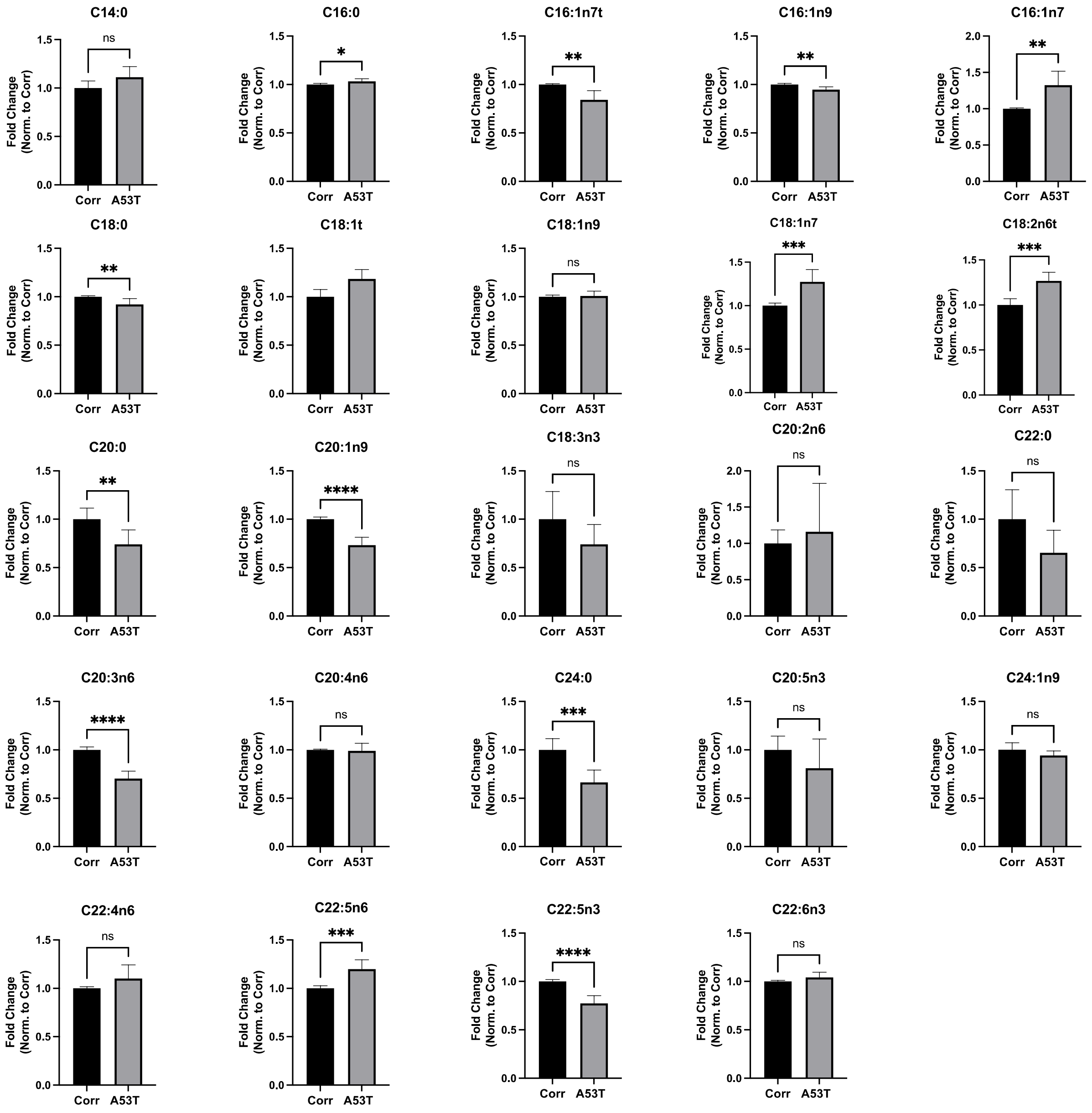


**Supplementary Figure 4.** Fatty acid profiles of the A53T and the Corr neurospheres at day 54.

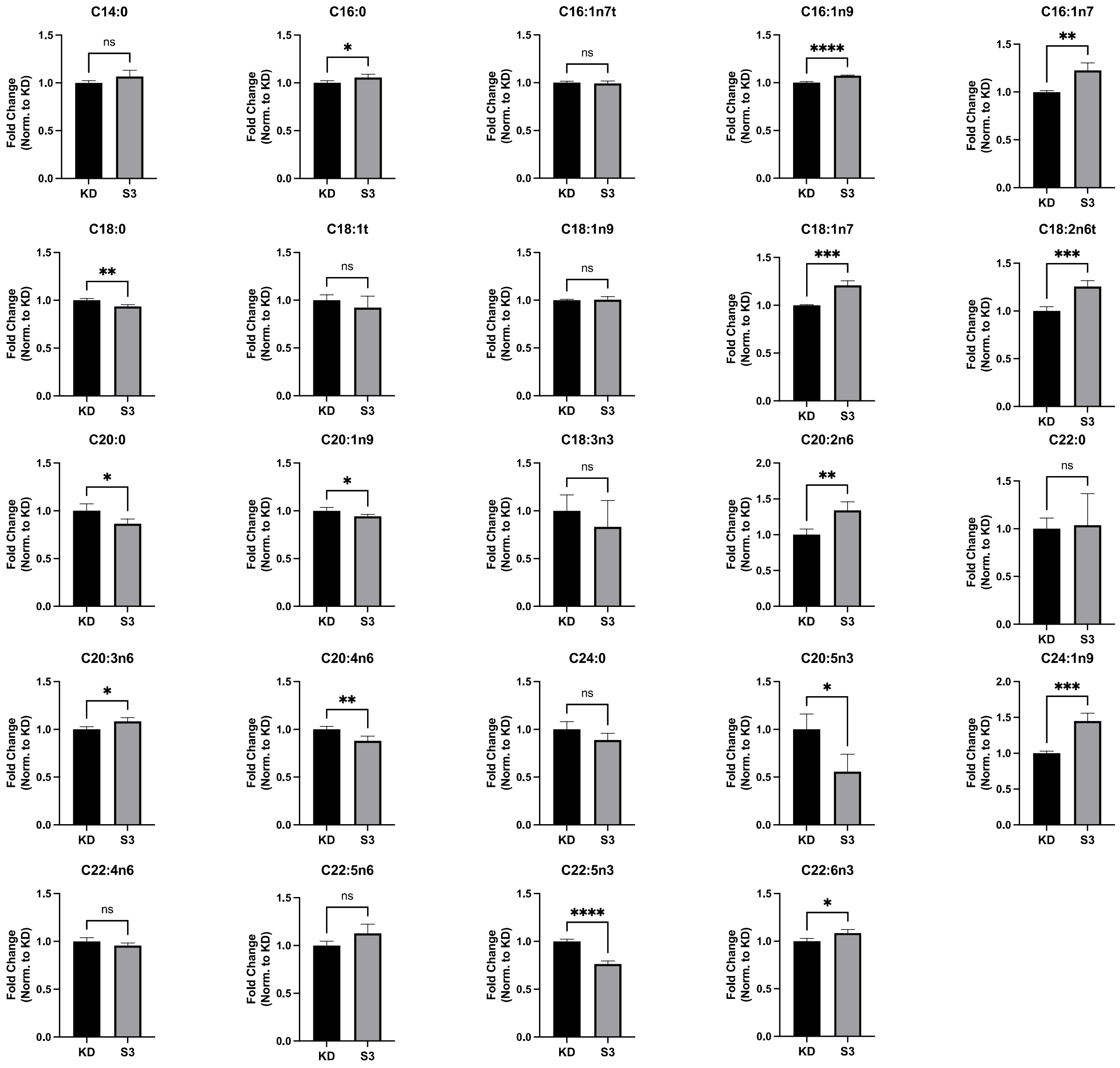


**Supplementary Figure 5**. Fatty acid profiles of the S3 and the KD neurospheres at day 54.

**Media and Supplements**

| Cell line | Media | Supplier | Catalog # | Supplement | Supplier | Volume (mL) | Catalog # |
|---|---|---|---|---|---|---|---|
| **iPSC** | mTesR1 (400 mL) | SCT | 85851 | mTesR1 Supplement | SCT | 100 | 85852 |
| **NSC Generation** | NeuroBasal (490 mL) | TFS | 21103049 | Neural Induction supplement | TFS | 10 | A1647701 |
| **NSC Expansion/ Neurosphere Plating** | NeuroBasal (245 mL) | TFS | 21103049 | Neural Induction supplement | TFS | 10 | A1647701 |
| | Advanced DMEM/F12 (245 mL) | | 12634028 | | | | |
| **Neurosphere Differentiation** | BrainPhys (470 mL) | SCT | 05790 | B27 | TFS | 10 | 17504044 |
| | | | | N2 | TFS | 5 | 17502048 |
| | | | | NEAA | TFS | 5 | 11140050 |
| | | | | GlutaMAX | TFS | 5 | 35050061 |
| | | | | PS | TFS | 5 | 15140122 |
| | | | | BDNF (500x) | Pepro Tech | 1 | 450-02 |
| | | | | GDNF (500x) | Pepro Tech | 1 | 450-10 |
| | | | | dbCAMP (500x) | Sigma-Aldrich | 1 | D0260 |
| **Neurosphere Maturation** | iCell Neural Base Medium 1 (480 mL) | CDI | M1010 | iCell Neural Supplement B | CDI | 10 | M1031 |
| | | | | iCell Nervous System Supplement | CDI | 5 | M1029 |
| | | | | PS | TFS | 5 | 15140122 |

**Abbreviation:- StemCell Technology (CST), Thermo Fischer Scientific (TFS), Cellular Dynamics International Inc.(CDI), Brain-derived neurotrophic factor (BDNF), Glial cell-derived neurotrophic factor (GDNF), Penicillin-Streptomycin (PS), Non-Essential Amino Acids (NEAA), N6,2′-O-Dibutyryladenosine 3′,5′-cyclic monophosphate sodium salt (dbCAMP)**

**Supplementary Table 1. Media composition and information.** All media and supplements used in the generation/maintenance of NSC and neurosphere are shown.

| Primary Antibody | Catalog # | Company | Host | ICC Dilution | Western Dilution |
|---|---|---|---|---|---|
| **Anti-MAP2** | 822501 | BioLegend | Chicken | 1:1000 | |
| **Anti-PAX6** | 426600 | Invitrogen | Rabbit | 1:200 | |
| **Anti-S100B** | S2532 | Sigma-Aldrich | Mouse | 1:500 | |
| **Anti-TAU** | A0024 | Dako | Rabbit | 1:500 | |
| **Anti-VGLUT2** | 135 403 | Synaptic Systems | Rabbit | 1:500 | |
| **Anti-GAD1/67** | MAB5406 | Millipore Sigma | Mouse | 1:500 | |
| **Anti-a-Synuclein** | 610787 | BD Biosciences | Mouse | | 1:1000 |
| **Anti-pS129 a-Synuclein** | 168381 | Abcam | Rabbit | | 1:500 |
| **Anti-Vinculin** | 13901 | Cell Signaling Technologies | Rabbit | | 1:1000 |
| **Anti-GAPDH** | ab8245 | Abcam | Mouse | | 1:2000 |

| Secondary Antibody | Catalog # | Company | ICC Dilution | Western Dilution |
|---|---|---|---|---|
| **Goat anti-Rabbit Alexa Fluor 488** | A11034 | Invitrogen | 1:500 | |
| **Goat anti-Chicken Alexa Fluor 555** | A21437 | Invitrogen | 1:500 | |
| **Goat anti-Mouse Alexa Fluor 647** | A21236 | Invitrogen | 1:500 | |
| **IRDye 800CW – Dk α Ms** | 926-32212 | LiCor | | 1:5000 |
| **IRDye 800CW – Dk α Rb** | 926-32213 | LiCor | | 1:5000 |
| **IRDye 680RD – Dk α Ms** | 926-68072 | LiCor | | 1:10,000 |
| **IRDye 680RD – Dk α Rb** | 926-68073 | LiCor | | 1:10,000 |

**Supplementary Table 2.** List of primary and secondary antibodies used for Immunostaining and Western blotting

| Gene | Species | Assay ID | Reporter | Catalog # |
|---|---|---|---|---|
| **MAP2 (microtubule associated protein 2)** | Human | Hs00258900_m1 | FAM | 4331182 |
| **PAX6 (paired box 6)** | Human | Hs01088114_m1 | FAM | 4331182 |
| **Synapsin I** | Human | Hs00199577_m1 | FAM | 4331182 |
| **S100B (S100 calcium binding protein B)** | Human | Hs00902901_m1 | FAM | 4331182 |
| **GAPDH (glyceraldehyde-3-phosphate dehydrogenase)** | Human | Hs03929097_g1 | VIC | 4448490 |

**Supplementary Table 3:** List of TaqMan Probes used for quantitative real-time PCR.

### Supplementary Materials and Methods

#### iPSC generation, Editing and Maintenance

All cells were maintained in the incubator at 37°C with 5% $CO_2$. The A53T patient-derived iPSC were generated from fibroblasts obtained from a patient skin biopsy at Boston University. The fibroblast to iPSC reprogramming was performed at the Harvard stem cell core using a non-integrative (mRNA) strategy. The α-syn triplication patient cell line (S3) was purchased from the Coriell institute for medical research. The iPSC were expanded and maintained in mTESR media from stem cell technology (see iPSC Media in **Supplementary Table 1**).

The disease-causing A53T mutation was corrected in house using the CRISPR-Cas9 system to generate an isogenic control pair. Multiple clones were evaluated and confirmed to have a normal karyotype. Similarly, the S3 mutation was corrected in house using the same method to generate an isogenic control for the S3 iPSC. The disease-causing A53T mutation was corrected in house using the CRISPR-Cas9 system. Briefly, guide RNAs were designed using the Zhang Lab Guide Design Tool (http://crispr.mit.edu) and cloned into pSpCas9(BB)-2A-GFP (PX458) (Addgene, Plasmid #48138). All clones were Sanger sequenced to verify inserted gRNA sequences. gRNA cleavage efficiencies were tested in 293T cells and analyzed by TIDE software (http://shinyapps.datacurators.nl/tide/) Guide 5' GTGGTGCATGGTGTGGCAAC 3' was used for A53T correction. A 127 nt ssODN with sequence 5' ATTTCATAGGAATCTTGAATACTGGGCCACACTAATCACTAGATACTTTAAATATCA TCTTTGGATATAAGCACAATGGAGCTTATCTGTTGCCACACCATGCACCACTCCCT CCTTGGTTTTGGAG 3' were synthesized at IDT and used as correction template. 2 ug pX458 plasmid and 1 ug ssODN was transfected to 1x $10^6$ A53T patient iPSCs using Lipofectamine 3000 (Cat. No. L3000008) and sorted for GFP positivity. Sorted cells were plated on Matrigel coated surface at 150 cell/$cm^2$ to allow the formation of single-cell derived clones. After 7-10 days of culture, colonies were picked into 96 w/p, for expansion and genotyping. Genomic DNA of each clone was prepared using Zygem prepGEM Universal kit (VWR Cat. No. 76218-666). Targeted region was PCR amplified using Forward primer 5' GCCCCGGTGTTATCTCATTCT and Reverse primer 5' TGTCCAAGGGTGTTTCCTGA. PCR products were Sanger sequenced using sequencing primer 5' GCAGTTTGTCAATACATTTTTGG. Clones with A53T correction along with a few non-targeted clones were expanded, banked and karyotyped.

Similarly, a-Syn triplication S3 line were targeted with tested gRNA 5' GTAAAGGAATTCATTAGCCA. Targeted region was PCR amplified using Forward primer 5' TAGCCAAGATGGATGGGAGATG and Reverse primer 5' CCATCACTCATGAACAAGCACC. PCR products were Sanger sequenced using sequencing primer 5' AGAGTCTCACACTTTGGAGGGT. Sequencing results were analyzed by TIDE software (http://shinyapps.datacurators.nl/tide/). Clones with indels were further TOPO cloned to resolve the allelic editing event. Final clones with two copy SNCA deletion, resulting from frame-shift causing indels, were expanded, banked and karyotyped.

#### Generation of Neural Stem Cells (NSC)

On day zero of differentiation, the iPSC (both disease and corrected isogenic control) were lifted from the surface as a single cell suspension using Accutase. After resuspending the iPSC, the Accutase was diluted using DPBS and the suspension was centrifuged to a obtain the cell pellet. This DPBS Accutase dilution process is referred to as “Accutase and DPBS” in this manuscript. The supernatant was aspirated, and the iPSC were re-plated onto a Matrigel coated 6 well plate in NSC Generation Media (see **Supplementary Table 1**) with Rock Inhibitor. On day one, the culture should be approximately 20% confluent. The media was replaced with NSC Generation Media without Rock inhibitor, followed by replenishing the media every other day until day seven. Any non-neural or unwanted colonies were manually scraped away. On day seven of differentiation, the cells were suspended into a single cell suspension using Accutase and DPBS, and re-plated at a density of $2 \times 10^5$ cells/cm$^2$ on a Matrigel coated 6 well plate in NSC Generation Media with Rock inhibitor. The following day, the media was switched to NSC Expansion Media (NEM, see **Supplementary Table 1**). The NSC were examined under the microscope for unwanted neural crest cells (larger and flatter cells). The neural crest cells were removed by Accutase treatment. Briefly, the cells were washed with DPBS, incubated in Accutase for two minutes at 37$^o$C, then transferred to a microscope at room temperature. The cells were observed under the microscope and kept in the Accutase until all the flat cells have lifted off the plate. Once all the neural crest cells had lifted, the Accutase was aspirated, and the culture was gently rinsed with DPBS three times. The remaining cells are suspended into a single cell suspension in NEM with Rock inhibitor and re-plated at $2 \times 10^5$ cells/cm$^2$ on a Matrigel coated plate. The following day, the media is replaced with fresh NEM without Rock inhibitor, and the cultures were monitored for neural crest cells under the microscope for the next few days. If neural crest cells were observed, the Accutase cleaning step was performed as described above. After achieving a pure population of NSC, the NSC were transferred onto poly-L-ornithine/laminin or poly-D-lysine/laminin coated surfaces. The NSC were expanded and frozen in Synth-a-freeze Cryopreservation Medium (**Figure 1A**).

#### Differentiation of NSC into neurons in 2D

NSC were maintained and expanded in NEM. For 2D neuron differentiation, the NSC were plated on a poly-D-lysine/laminin coated surface in NEM. The next day, neuron differentiation media with CultureOne supplement was added to the NSC (the same as neurosphere Differentiation Media in **Supplementary Table 1**). The cells were kept in this media for four days, with a daily media change. On day 4, the neurons were re-plated onto a new poly-D-lysine/laminin coated surface in the desired plate format. The neurons were aged for 2 to 4 weeks in neuron maturation media (the same as neurosphere Maturation Media in **Supplementary Table 1**). At the end of the experiment, the neurons were washed once with DPBS and stored in -80°C (for RNA or protein analysis) or fixed in 4% paraformaldehyde for immunocytochemistry.

#### Generation of neurospheres

To plate the NSC for neurosphere differentiation, NSC were lifted from the surface as a single cell suspension using Accutase. After washing out Accutase, the NSC were resuspended in NEM and strained through a 40 micron cell strainer. The density of the cell suspension was adjusted to plate 20,000 cells per well. The cell suspension was added to the 384 well ultra-low adhesion spheroid plates, leaving the outermost rows and columns filled with sterile water. After loading cells, the spheroids plates were centrifuged at 300g for 5 minutes to aggregate the cells in the bottom of the well and to remove any air bubbles in the media. The NSC formed spheroids overnight in each well. The next day, around 80% of the NEM was aspirated and replaced with the neurosphere Differentiation Media (see **Supplementary Table 1**) to initiate differentiation. At this point, CultureOne can be added to the media to increase the neuronal purity of the cultures. The differentiation media was changed every two or three days. After ten days of differentiation, the neurosphere Differentiation Media was replaced by the neurosphere Maturation Media (see **Supplementary Table 1**). The media was changed bi-weekly for the next three weeks, and then once a week after day 30. The total media volume was approximately 90 µl per well in the 384 well plates. In each media change, 60 µl was aspirated and same volume was added back to each well. The neurospheres were harvested at different time points using wide orifice pipette tips for characterization. The desired number of spheres were added to microfuge tubes, washed with DPBS and stored in -80°C (for RNA, protein, or FADI analysis) or fixed in 4% paraformaldehyde for cryosectioning (**Figure 1A**).

#### Compound treatment

40 day-old neurospheres from both patient-derived and isogenic control lines were treated with 0.3 uM CAY10566 along with DMSO as a vehicle. The final concentration of DMSO in the media was 0.03%. The spheres were treated twice a week and harvested after two weeks of treatment. The spheres were harvested and pooled from multiples wells of the 384 well plates for different assays. The spheres were harvested and washed once with the DPBS and processed for the appropriate assay.

#### RT-qPCR

NSC, neurons, and neurospheres were washed once with DPBS and processed for RNA extraction using the RNeasy Plus Mini Kit (from Qiagen). The amount of RNA was quantified using Nanodrop (Thermofisher). The extracted RNA was converted to cDNA with the qScript cDNA Supermix (from Quanta Biosciences) and the Mastercycler ep using RT-PCR. The cDNA was loaded in a 96 well or 384 well qPCR plate at a starting RNA concentration of 20 ng/ul for all the samples. Gene expression was then analyzed with qPCR using the Taqman Fast Advanced Master Mix (from Thermofisher), Taqman Gene Expression Probes, and the Step One Plus Real-Time PCR System. The list of genes run for qPCR are PAX6, Nestin, SOX10, MAP2, S100B, VGLUT2, Synapsin I, and GAD2, and normalized with the housekeeping gene GAPDH. The catalog number of all the TaqMan probes are listed in **Supplementary Table 4.**

#### Immunocytochemistry

NSC and neurons were cultured in 24 or 96 well plates, washed once with DPBS, fixed for 30 minutes in 4% paraformaldehyde (PFA), and stored at 4°C in DPBS with 0.05% sodium azide. NSC were stained for different neuronal stem cell markers (PAX6 and Nestin), as well as the neural crest marker SOX10. Neurons were stained for different neuronal markers (MAP2, total Tau, VGLUT2 and GABA) as well with the astrocyte markers S100B.  neurospheres were harvested as mentioned above and fixed in 4% PFA overnight. The PFA was washed out and neurospheres were stored at 4° C in PBS with 0.05% sodium azide. The neurospheres were cryosectioned into 14 µm sections on glass slides (by Histoserv Inc). The sections were stained using a published protocol (Raja et al., 2016). Briefly, the sections were washed once with PBS and permeabilized for 30 minutes in PBS containing 0.3% Triton-X100 (PBST) and blocked in PBST with 10% v/v normal Goat (or Horse) serum for 1 hour. The blocking solution was replaced with primary antibody solution and incubated overnight at 4°C in PBST containing 5% normal Goat/Horse serum. Sections then received three 15 min washes in PBST containing 5% normal Goat/Horse serum. Secondary antibodies were prepared in PBST containing 5% Goat/Horse serum and Hoechst (for nuclear immunoreactivity) and sections were incubated in the secondary antibody solution for 2 hours at room temperature. The sections were washed three times in PBST containing 5% normal Goat/Horse serum, as described before. Sections were then cover slipped in Fluoromount-G mounting medium and the edges were sealed with nail polish. Slides were imaged on a Nikon confocal microscope. The catalog number and sources of all the primary and secondary antibodies are listed in **Supplementary Table 2 and 3.**

#### Western blotting

Approximately 60 to 80 neurospheres (or NSC pellets of 4-10 million cells) were harvested and stored at -80°C as mentioned above. The neurospheres were homogenized in lysis buffer (20 mM HEPES, 150 mM NaCl, 10% Glycerol, 1 mM EGTA, 1.5 mM $MgCl_2$, 1% Triton X-100, protease (Sigma P8340) and phosphatase (P2850, P5726) inhibitors) using a pestle. After homogenization, the samples were incubated on ice for 20 minutes and subjected to two freeze-thaw cycles. Following the freeze-thaw, samples were centrifuged at 14,000g for 30 minutes at $4^oC$ to sediment insoluble material.

Supernatant protein was harvested and measured using the BCA method (Thermofisher 23225). 40 ug of total protein was separated on 4-12% SDS PAGE gels (Invitrogen WG1401A), and dry transferred to 0.22 uM PVDF membranes. The membrane was fixed in 1% paraformaldehyde for 40 minutes prior to blocking for 30 minutes at room temperature (LiCor Odessey blocking buffer 1:1 in TBS). The membrane was incubated in primary antibodies overnight at 4°C with gentle rocking. Following primary antibody incubation, blots were washed three times for 15 minutes each in TBS solution containing 0.01% Tween20. The membrane was then incubated in secondary antibodies at room temperature for 2 hours, followed by three washes in 0.01% tween20 in TBS solution. After the washes, the blots were imaged on a Li-Cor Odyssey CLX and analyzed on LiCor ImageStudio software. The catalog number and sources of all the primary and secondary antibodies are listed in **Supplementary Table 2 and 3.**

#### Fatty Acid Desaturation Index (FADI) Analysis

The neurospheres were harvested as mentioned above, washed once with DPBS and resuspended in 350 µl cold 80% methanol and stored at -80°C until ready to process. The samples were shipped (on dry ice) to OmegaQuant LLC and processed for different fatty acid using gas chromatography (GC) with flame ionization detection. The neurosphere solution (80% methanol) was transferred into a screw-cap glass vial and dried in a speed vac. After drying, methanol containing 14% boron trifluoride were added and the vials were briefly vortexed and heated in a hot bath at 100°C for 10 minutes. After cooling, hexane and HPLC grade water were added sequentially. The vials were recapped, vortexed and centrifuged to separate layers. An aliquot of the hexane layer was transferred to a GC vial and processed. Fatty acid composition was expressed as a percent of total identified fatty acids. The abundance of different fatty acid was quantified using Microsoft Excel and plotted using GraphPad Prism software.

#### Statistical Analysis

The results are presented as the mean ± standard deviation. All statistical analyses were performed using GraphPad Prism 7.0 software. The p values were calculated with a two-tailed Student's t-test and One way ANOVA (Tukey's test $*p < 0.05$, $**p < 0.01$, $***p < 0.001$). Results shown here are the representative data from several experiments performed independently at different time points by multiple individuals.
